# Proactive visual and motor prioritization differentially scale with cue reliability

**DOI:** 10.64898/2026.01.28.702371

**Authors:** Sisi Wang, Freek van Ede

## Abstract

Environmental cues enable the brain to anticipate and prepare for upcoming behavior, such as by selectively prioritizing relevant visual representations and associated action plans in working memory in service of an imminent task. While it has been demonstrated that neural dynamics of visual and motor prioritization each scale with cue reliability, studies to date tracked either visual or motor prioritization in isolation. It therefore remains unknown whether visual and motor prioritization scale similarly or differently with cue reliability. To fill this gap, we manipulated cue reliability (100%, 80%, 60%) in a visual-motor working-memory task that uniquely enabled us to isolate the neural dynamics associated with visual and motor prioritization in anticipation of an imminent working-memory task. EEG measurements in male and female human volunteers revealed how cue reliability differentially drives visual and motor prioritization. While the strength and timing of visual prioritization were relatively stable across cue reliability levels, motor prioritization profoundly scaled with cue reliability and developed more gradually with lower certainty. These findings show that visual and motor prioritization in working memory are differentially susceptible to the certainty conveyed by environmental cues, and suggest that motor prioritization may be more cautious in nature.

**Significance Statement:** To cope with the ever-changing world, the human brain continuously leverages environmental cues to anticipate upcoming behavior, such as by prioritizing relevant visual representations and action plans ‘in mind’. Yet, in a volatile world, environmental cues typically vary in the certainty they provide. Building on prior work studying visual or action prioritization in isolation, we uniquely studied how cue certainty shapes both visual and motor prioritization within the same task. We unveil how cue certainty distinctly drives visual and action prioritization, with action prioritization requiring more certainty before deployment, whilst also being deployed more gradually at lower certainty. Thus, prioritization of potential actions is distinct from—and more cautious than—prioritization of the visual representations that guide these actions.

## Introduction

The brain is proactive, continuously picking up and leveraging cues from the external environment to prepare for upcoming behavior. For example, cues conveying what working-memory representations are most likely to become relevant for an imminent task drive the selective prioritization of specific visual representations and their associated action plans (e.g., Schneider et al., 2017; van Ede et al., 2019; Boettcher et al., 2021; Nasrawi et al., 2023, 2025; Echeverria-Altuna et al., 2025; Gresch et al., 2025). Yet, in the dynamic volatile world we face in everyday life, environmental cues rarely provide full certainty, but vary in the degree of certainty they provide.

It has been well established that cue reliability shapes anticipatory neural dynamics associated with preparation for an upcoming task. Cue reliability has been shown to modulate the spatial lateralization of 8-12 Hz alpha activity associated with task anticipation, when anticipating sensory input (Gould et al., 2011; Haegens et al., 2012), or prioritizing visual representations in working memory for an imminent task (Günseli et al., 2019; Fu et al., 2022; Wang and van Ede, 2025a). In a parallel line of investigation, cue reliability has also been shown to modulate EEG mu/beta dynamics associated with planning for a potential action (e.g., Tzagarakis et al., 2010, 2015, 2021).

While cue reliability has thus been shown to shape both sensory and motor (action) prioritization, studies to date exclusively considered sensory or motor prioritization in isolation. Accordingly, it remains unclear whether cue reliability similarly or differentially modulates sensory and motor prioritization (and comparing findings in existing literature is challenging because studies used different tasks and participants). Knowing how visual and motor prioritization are jointly susceptible to cue reliability is relevant because sensory representations often serve to guide specific actions (Heuer et al., 2020; Olivers and Roelfsema, 2020; van Ede, 2020). Moreover, while it has been shown how visual and motor prioritization in working memory can co-develop (van Ede et al., 2019; Gresch et al., 2025), recent evidence shows that visual and motor prioritization can also de-couple (Echeverria-Altuna et al., 2025; Nasrawi et al., 2025). Whether visual and motor prioritization co-develop or de-couple under varying levels of cue reliability remains unaddressed.

To fill this gap, we build on a recently developed visual-motor working-memory task (as in van Ede et al., 2019; Boettcher et al., 2021; Nasrawi et al., 2023, 2025; Echeverria-Altuna et al., 2025; Gresch et al., 2025) that enabled us to independently track visual and motor prioritization in the same task, in the same participants. Participants encoded two visual objects in working memory that were linked to two potential manual action via their tilt. Following a central color-cue – that was 100%, 80%, or 60% reliable – we could study the selective prioritization of the cued visual representation, and its associated manual action. We did so through two established EEG markers: posterior 8-12 Hz alpha lateralization according to memorized item location (e.g., Myers et al., 2014; Poch et al., 2014; Wallis et al., 2015; Mok et al., 2016; Schneider et al., 2017; van Ede et al., 2017, 2019; Liu et al., 2022; Woodman et al., 2022; Wang and van Ede, 2024) and central 8-30 Hz mu/beta lateralization according to the associated response hand (e.g., Kaiser et al., 2003; Pfurtscheller et al., 2005; Schneider et al., 2017; van Ede et al., 2019). Because the cued object’s location (visual attribute) and prospective response hand (motor attribute) were orthogonally manipulated, we could independently track EEG markers of visual and motor prioritization, and study them as a function of cue reliability.

To preview our results, we report that cue reliability differentially modulates our neural markers of proactive visual and motor prioritization in anticipation of an imminent task. While the strength and timing of our visual-prioritization marker remained relatively stable to cue reliability levels, our motor-prioritization marker profoundly scaled with cue reliability, developed more gradually with lower certainty, and became undetectable following 60%-reliable cues (despite robust visual prioritization in the same condition).

## Methods

### Participants

Twenty-five healthy human volunteers participated in the experiment (age range from 18 to 32; mean age = 21.9; SD = 3.4; 18 females and 7 males). Sample size was set a-priori to n = 25 based on previous studies from our lab that had similar designs and focused on similar EEG outcome variables (van Ede et al., 2019; Boettcher et al., 2021; Nasrawi et al., 2023, 2025; Echeverria-Altuna et al., 2025; Gresch et al., 2025; Wang and van Ede, 2025a, 2025b). One participant was excluded from EEG data analyses due to large artifacts in their EEG signal (less than 70% artifact-free trials remaining). The experimental procedures were reviewed and approved by the Research Ethics Committee of the Faculty of Behavioral and Movement sciences Vrije Universiteit Amsterdam. All participants provided written informed consent prior to participation and were reimbursed 10 Euros per hour as a compensation for their time.

### Task and Procedure

Human participants performed a visual–motor working-memory task in which color cues (retrocues; Griffin and Nobre, 2003; Souza and Oberauer, 2016; Van Ede and Nobre, 2023) presented during the maintenance interval indicated which of two memorized items—a pair of differently colored, oriented gratings—was most likely to be tested after a second delay (**Fig. 1a**). Across blocks, cue reliability was manipulated such that cues were 100%, 80%, or 60% predictive of the to-be-tested memory item (**Fig. 1b**).

**Figure 1.**
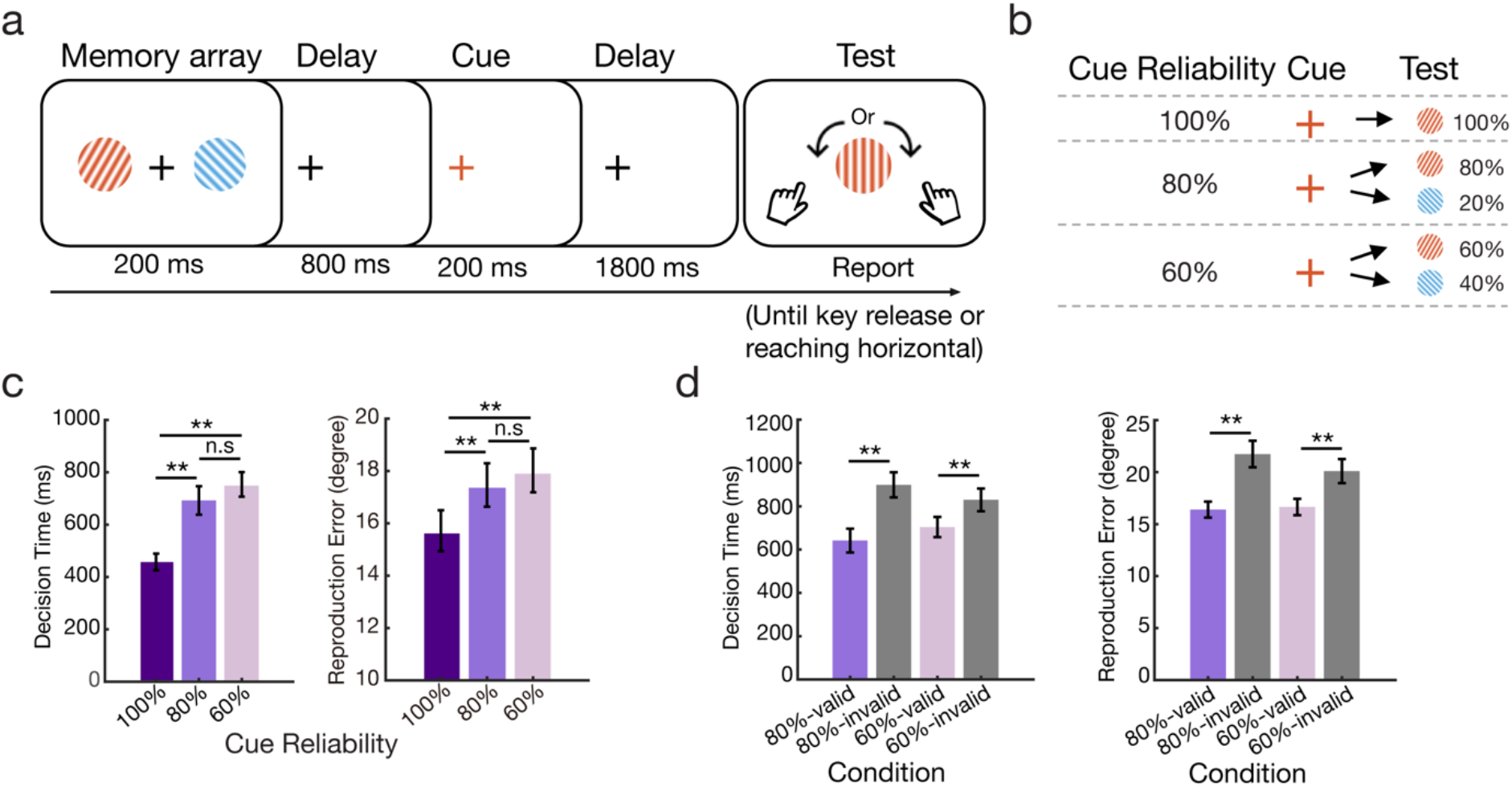
Task paradigm and behavioral performance. **a)** The experimental procedure of the visual-motor working-memory task. **b)** Illustration of different cue reliability conditions. **c)** Participants’ (n = 25) average behavioral performance across three levels of cue reliability; left panel: decision time, right panel: reproduction error. **d)** Participants’ average behavioral performance across different cue reliability (100%, 80%, and 60% reliable cues) and cue validity (valid and invalid cue trials) conditions; left panel: decision time, right panel: reproduction error. Error bars represent ± 1 SEM. *, **, n.s represent significance level p<0.05, p<0.01, and non-significant after Bonferroni correction, respectively.

Our central aim was to assess how prospective certainty about the upcoming working-memory test influenced the proactive prioritization of the cued visual item and its associated action plan following the predictive cue. To this end, we tracked two established EEG markers of visual and motor prioritization: (1) lateralization of posterior 8–12 Hz alpha activity according to the cued item’s memorized location indexing visual prioritization, and (2) lateralization of central 8–30 Hz mu/beta activity according to the cued item’s prospective response hand indexing motor prioritization. Critically, we orthogonally manipulated the spatial location of each item (left/right) and its associated response hand (left/right tilt mapped to left/right prospective keypress), enabling independent tracking of visual prioritization (prioritization of the item’s location) and motor prioritization (prioritization of the prospective response hand). Building on several recent studies that employed this same approach for independently tracking visual and motor prioritization in working memory (van Ede et al., 2019; Boettcher et al., 2021; Nasrawi et al., 2023, 2025; Echeverria-Altuna et al., 2025; Gresch et al., 2025), we here uniquely added a manipulation of cue reliability to assess how proactive visual and motor prioritization processes each scale with certainty of the anticipated working-memory test.

Each trial began with the presentation of two colored, tilted gratings for 200 ms, followed by an 800 ms delay where only the central fixation cross remained in the screen. The fixation cross then changed color for 200 ms to match one of the memorized gratings, serving as the cue. After a further 1800 ms delay, participants were shown a vertically oriented probe stimulus (0°) at the center of the screen, and were required to reproduce the memorized orientation of the memory item that matched the color of this probe stimulus. They did so by pressing a response key with the corresponding hand (“Z” for left-tilted responses, “?/” for right-tilted responses). Upon key press, the dial would tilt (in a speed of 120°/sec) in the left/right direction until the key was released, terminating the response. It was not possible to adjust the dial after key release, and the dial could be rotated by a maximum of 90°. Because of these features, a left (right) tilted memory item could only be accurately reported by pressing the left (right) button and holding it down until reaching the desired tilt. After each response, a score from 0-100 indicated reproduction accuracy (100 = perfect reproduction; 0 = 90° error).

Cue reliability was manipulated in separate blocks. In the 100% condition, the cue always matched the color of the subsequently probe, and thus informed with 100% reliability which memory item would become tested. In the 80% condition, the cue was valid on 80% of trials and invalid on 20%; in the 60% condition, the cue was valid on 60% and invalid on 40%. The color of the probe stimulus at the memory test always indicated which memory item to report.

Each participant completed two experimental sessions, each containing all three cue-reliability blocks (100%, 80%, 60%), with block order pseudorandomized and counterbalanced across participants. Before the first session, participants practiced 20 trials of each reliability condition to familiarize themselves with the task. They continued to the formal experiment once practice accuracy exceeded 75% (~22.5° mean error). In each session, participants completed three blocks of 160 trials (one block per cue-reliability condition), yielding a total of 960 trials across the experiment (~2.5 hours).

### Apparatus and stimuli

Stimuli were presented using MATLAB (R2020a; MathWorks) and the Psychophysics Toolbox (version 3.0.16; Brainard, 1997) on a 23-inch LED monitor (1920 × 1080 resolution; 240 Hz refresh rate). Participants were seated 70 cm from the screen with their head stabilized using a chin rest to ensure consistent viewing and stable EEG recording.

Throughout the experiment, the display contained a gray background (RGB: 128, 128, 128) and a central fixation cross (0.75°). Memory and test stimuli were high-contrast sine-wave gratings (100% contrast; 4.6° diameter; spatial frequency: 0.032 cycles/pixel; phase: 90°). During encoding, two gratings appeared 6° to the left and right of fixation. To maintain the orthogonal mapping between item location and response hand, each memory array contained one left-tilted (−80° to −10°) and one right-tilted (10° to 80°) grating. One grating was colored blue (RGB: 21, 165, 234) and the other orange (RGB: 234, 74, 21). The mapping between item color, location, and orientation were randomized across trials. For example, trials in which the left memory item was cued, this item was equally often a left-tilted or a right-tilted grating. This ensured that item location (our key visual attribute) and prospective response hand linked to item tilt (our key motor attribute) were orthogonal across trials, enabling independent tracking of visual and motor prioritization following the cue.

Unlike the two lateralized memory items, the memory test stimulus (probe) at the end of each trial always appeared centrally. It was always the same size as the memory gratings at encoding. Its initial orientation was always vertical (0°) and its color (blue or orange) indicated which item to report and therefore which response hand was required. Probe color matched the cued item’s color on valid-cue trials and the other item’s color on invalid-cue trials.

### Analysis of behavioral data

Decision time (ms) was defined as the interval between the onset of the probe stimulus (the vertically oriented central test stimulus) and the initiation of the participant’s response. Reproduction error (degrees) was quantified as the absolute angular difference between the reported orientation and the true orientation of the tested item. Trials with decision times exceeding 4 seconds were excluded from all analyses.

### EEG acquisition and pre-processing

EEG was recorded using a 64-channel BioSemi ActiveTwo system (1024 Hz sampling rate), with electrodes arranged according to the international 10–20 system. The CMS and DRL electrodes, positioned near POz, served as the online reference, and data were re-referenced offline to the average of the left and right mastoids. EOG electrodes were placed around the eyes and were used exclusively for artifact removal via independent component analysis (ICA).

Offline preprocessing was performed in MATLAB using FieldTrip (Oostenveld et al., 2011) and custom scripts. After re-referencing, the continuous EEG was segmented from −200 to 2000 ms relative to cue onset. Fast ICA (implemented in FieldTrip) was applied to remove components reflecting eye blinks and horizontal eye movements, identified through correlations with the VEOG (vertical EOG difference) and HEOG (horizontal EOG difference) signals. Remaining artifacts were removed through visual inspection using FieldTrip’s *ft_rejectvisual* function with the “summary” method. Trial exclusions were performed on all trials, without knowledge of trial-specific condition labels. We excluded participants with excessive artifacts in their EEG data (i.e., fewer than 70% artifact-free trials remaining) to ensure sufficient data available for our EEG data analysis. One participant was excluded due to excessive artifact removal (only 69.8% usable trials remaining). For the remaining participants, an average of 93.0% of trials (SD = 4.3%) were retained (out of 320 trials per condition), corresponding to mean counts of 297.3, 296.2, and 299.0 artifact-free trials for the 100%, 80%, and 60% cue reliability blocks, respectively.

### EEG time-frequency analysis

For the EEG time–frequency decomposition, we first applied a surface Laplacian transform to enhance the spatial specificity of the EEG signal (Babiloni et al., 2001; Carvalhaes and De Barros, 2015; Kayser and Tenke, 2015). Clean EEG epochs were then decomposed into time–frequency representations using a short-time Fourier transform with Hanning tapering (FieldTrip’s *ft_freqanalysis*). Spectral power from 2–40 Hz (1-Hz steps) was estimated using a 500-ms sliding window advanced in 10-ms steps.

#### Posterior alpha lateralization

Alpha-band power (8–12 Hz) was extracted and used to compute lateralization on a trial-by-trial basis. Following prior work (van Ede et al., 2019; Boettcher et al., 2021; Nasrawi et al., 2023, 2025; Echeverria-Altuna et al., 2025; Gresch et al., 2025), lateralization was quantified at electrodes PO7/PO8 using the normalized asymmetry index: ((contralateral electrode – ipsilateral electrode) / (contralateral electrode + ipsilateral electrode)) × 100. For each trial, contralateral and ipsilateral electrodes were defined relative to the spatial location (left vs. right) of the cued memory item. Trial-wise values were then averaged to obtain the participant-level alpha lateralization time course.

#### Central mu/beta lateralization

The same procedure was applied to compute motor-related lateralization in the mu/beta band (8–30 Hz). Power was extracted from electrodes C3/C4 (as in van Ede et al., 2019; Boettcher et al., 2021; Nasrawi et al., 2023, 2025; Echeverria-Altuna et al., 2025; Gresch et al., 2025), and lateralization was computed using the same asymmetry formula, with contralaterality defined relative to the prospective response hand (left vs. right) of the cued memory item. Trial-wise indices were averaged to yield the overall mu/beta lateralization profile.

#### Topographical representations

To visualize lateralization across the scalp, we computed topographical maps by contrasting left versus right item location (for alpha) and left versus right response hand (for mu/beta) trials using the corresponding normalized contrast: (i.e., ((left − right) / (left + right)) × 100).

### Experimental Design and Statistical Analyses

The experiment is a within-subject design including three cue-reliability conditions (100%, 80%, and 60%). Behavioral performance was analyzed using one-way repeated-measures ANOVAs on decision times and absolute reproduction error across the three reliability levels. For conditions containing both valid and invalid trials (80% and 60% blocks), paired-samples t-tests were used to compare valid versus invalid trials. Multiple comparisons were corrected using the Bonferroni method, and all reported p values are Bonferroni-adjusted.

To evaluate the temporal dynamics of EEG lateralization (posterior alpha and central mu/beta), we used cluster-based permutation testing (Maris and Oostenveld, 2007) implemented in FieldTrip (*ft_timelockstatistics*, ‘montecarlo’ method). This approach controls for multiple comparisons by evaluating spatio-temporally adjacent data points under a single permutation-derived cluster distribution. We generated 10,000 permutations to estimate the null distribution of the largest cluster. Clusters were identified using FieldTrip’s default settings, grouping temporally adjacent points showing significant mass-univariate t-tests (two-sided α = 0.025). Cluster size was defined as the summed t values within each cluster. This procedure was applied to test visual alpha and motor mu/beta lateralization against zero and to compare these measures across cue-reliability conditions.

In addition, we estimated regression coefficients to assess how cue reliability predicted visual and motor prioritization. Trial-wise alpha- and mu/beta-band lateralization were first entered into a general linear model at the single-subject level, with cue reliability as a predictor at each time point. The resulting regression coefficients (β estimates) were then averaged across participants for alpha and mu/beta lateralization at each time point. These group-level regression time courses were subsequently evaluated using cluster-based permutation testing.

To compare onset latencies of alpha and mu/beta lateralization across cue reliability conditions, we used jackknife-based latency estimates of the first datapoint that reached 25% of the peak value (Smulders, 2010). Latencies were analyzed using paired-samples t-tests to compare visual versus motor prioritization within each reliability level.

Bayesian factor analyses were further conducted to quantify the null effect of cue reliability on alpha lateralization using open-source software JASP (version 0.18.3, Quintana and Williams, 2018).

### Data and code availability

All data and analysis scripts are publicly available on OSF: https://osf.io/hbdpr/.

## Results

Human participants performed a visual-motor working-memory task in which cues (that were presented during working-memory maintenance) indicated which of two memorized items (colored, oriented gratings) would be most likely to become tested after another working-memory delay (**Fig. 1a**). Across blocks, cues varied in reliability—being 100%, 80%, or 60% predictive of the to-be-tested memory item (**Fig. 1b**). This afforded proactive prioritization of the cued visual representation and its associated manual action plan.

Our primary aim was to investigate how prospective certainty of which memory item would become tested influences the respective prioritization of visual and motor working-memory representations in anticipation of ensuing memory-guided behavior. To this end, we orthogonally manipulated whether the cued item was left or right at encoding, and whether it required a left- or right-hand response by virtue of its left/right tilt. Building on prior studies using this orthogonal manipulation (see: van Ede et al., 2019; Boettcher et al., 2021; Nasrawi et al., 2023, 2025; Echeverria-Altuna et al., 2025; Gresch et al., 2025), this enabled us to independently track “visual prioritization” (operationalized as prioritization of the left/right memorized item) and “motor prioritization” (operationalized as selecting the left/right tilted item that was linked to a left/right-hand response), to assess how each are modulated by cue reliability.

We tracked visual and motor prioritization through two well-established EEG markers: the lateralization of posterior 8-12 Hz alpha activity according to the cued item’s memorized location (visual prioritization) and the lateralization of central 8-30 Hz mu/beta activity according to the cued item’s associated manual action (motor prioritization).

We first present behavioral results to confirm that participants utilized the cues presented during the working-memory delay. After establishing this, we then turn to our primary EEG analyses of proactive visual and motor prioritization during working memory and delineate how cue reliability differentially shapes both forms of prioritization.

### Working-memory performance scales with cue reliability

Before we dive into our primary EEG results, we first evaluated how behavioral performance— decision time and reproduction error—was influenced by cue reliability. Decision time (in ms) was defined as the time from test onset (i.e., the vertical-oriented probe grating presented in the center) to response initiation (starting to ‘dial’ the probe grating left- or right-wards by holding down the left/right response key). Reproduction error (in degrees) was defined as the absolute value of the difference in orientation between the tested item’s orientation and the reported orientation.

We first evaluated decision times and reproduction errors across the blocks in which the cues were 100%, 80%, or 60% reliable for the item that would be tested (**Fig. 1c**). We observed a main effect of cue-reliability block type on both decision times and reproduction errors (decision time: *F*(2,48) = 42.700, *p* < 0.001, η_p_^2^ = 0.640; reproduction error: *F*(2,48) = 13.060, *p* < 0.001, η_p_^2^ = 0.352). Follow-up t-tests confirmed significantly faster decision times and lower reproduction errors in the 100% cue-reliable block compared to both the 80% and 60% blocks (decision time: |*t*|s ≥ 6.354, *p*s < 0.001; reproduction error: |*t*|s ≥ 4.118, *p*s ≤ 0.001, all p-values are Bonferroni-corrected). In contrast, we found no statistically significant difference between performance in the 80% and 60% cue-reliability blocks (decision time: *t*(24) = −1.932, *p* = 0.196; reproduction error: *t*(24) = −1.097, *p* = 0.850).

We next turned to the blocks with 80% and 60% reliable cues where we could additionally investigate the effect of cue validity: comparing trials where the color of the tested memory item was the same as the cued item (valid cue) or where the other item was tested instead (invalid cue). This comparison (**Fig. 1d**) confirmed better performance (faster decision time and smaller reproduction error) in valid-cue trials compared to invalid-cue trials, in both the 80% (decision time: *t*(24) = −7.729, *p* < 0.001; reproduction error: *t*(24) = −6.212, *p* < 0.001) and the 60% cue-reliability blocks (decision time: *t*(24) = −4.284, *p* < 0.001; reproduction error: *t*(24) = −3.792, *p* < 0.001).

Together, these results show that participants used the cues to prepare for the upcoming memory task, yielding the highest performance when cues were 100% reliable. When cues were less reliable, participants still used the cues as evidenced by better performance following valid vs. invalid cues in both the 80% and 60% cue-reliability conditions.

### Robust visual prioritization regardless of cue reliability

Having established robust cueing benefits on memory-guided performance, we next turned to our EEG marker of visual prioritization as reflected in posterior alpha lateralization according to the cued item’s memorized location.

As shown in **Figure 2a-c** (left panels), we observed clear cue-induced alpha lateralization (8–12 Hz) in all cue-reliability conditions, with a characteristic posterior topography (**Fig. 2a-c**, right panels).

**Figure 2.**
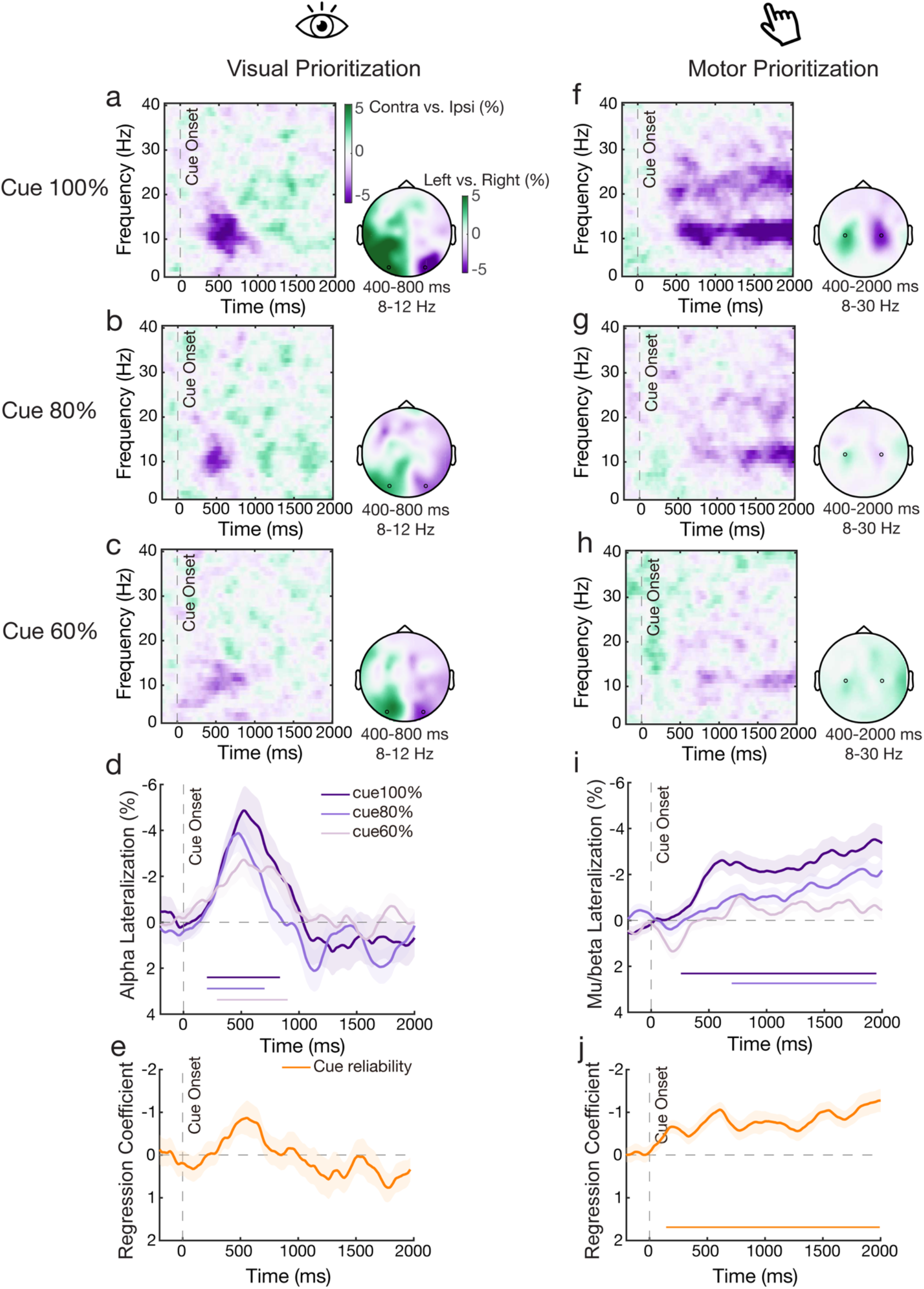
Cue reliability differentially modulates proactive visual and motor prioritization during working memory. **a)** Left panel: time-frequency spectrum of visual lateralization relative to the cued item’s memorized location in posterior electrodes (PO7/PO8, highlighted as the dark dots in the topographic plots) when the cue was 100% reliable. Right panel: topographic maps of alpha power (8-12 Hz) following 100% reliable cues averaged over 400-800 ms following cue onset. **b)** Same plots as in **a** following 80% reliable cues. **c)** Same plots as in **a** and **b**, following 60% reliable cues. **d)** Time courses of alpha lateralization (8-12 Hz) across three levels of cue reliability. **e)** The group-level average of beta estimates from general linear model fitting trial-wise alpha lateralization (8-12 Hz) with cue-reliability as a continuous predictor. **f-j)** Similar as **a-e**, but for 8-30 Hz motor lateralization in central electrodes (C3/C4, highlighted as the dark dots in the topographic plots) relative to the prospective response hand associated with the cued item. Colored horizontal lines above the x-axes indicate significant clusters (p<0.05). Shading and error bars represent ± 1 SEM, n = 24.

Comparisons of time courses of aggregated alpha lateralization (across 8-12 Hz over the posterior electrode pair PO7/PO8, as in Myers et al., 2014; Poch et al., 2014; Wallis et al., 2015; Mok et al., 2016; Schneider et al., 2016; van Ede et al., 2017, 2019; van Ede, 2018; Liu et al., 2022; Woodman et al., 2022; Wang and van Ede, 2024; across cue-reliabilities confirmed robust alpha lateralization in all cue-reliability blocks (**Fig. 2d**; 100% cluster-*p* < 0.001, 80% cluster-*p* < 0.001, 60% cluster-*p* < 0.001; comparable alpha lateralization results were observed when averaging over the posterior electrode cluster, PO7/8, PO3/4, P7/P8, P5/6, O1/O2; see **Fig. S1a-b**). Though alpha lateralization was numerically larger with larger cue-reliability (as also reported in Günseli et al., 2019; Fu et al., 2022; Wang and van Ede, 2024), direct comparisons between cue-reliability blocks did not yield significant differences (cluster-*p*s: 100% vs. 80%, *p* = 0.062; 100% vs. 60%, *p* = 0.116; 80% vs. 60%, *p* = 0.154). Further Bayesian factor analyses on the average amplitude of alpha lateralization (averaged over a pre-defined time window of 400–800 ms post-cue, consistent with van Ede et al., 2019; Wang and van Ede, 2025a) revealed only limited evidence in favor of the null hypothesis, particularly for comparisons between the 100% and 80% conditions, and between the 100% and 60% conditions (100% vs. 80%: BF_01_ = 1.44; 100% vs. 60%: BF_01_ = 0.70; 80% vs. 60%: BF_01_ = 4.52).

In addition to pairwise differences, we further ran a regression analysis to examine whether trial-wise alpha lateralization could be predicted by cue reliability. Consistent with aforementioned results, alpha lateralization could not be significantly predicted by cue reliability (**Fig. 2e**; no significant cluster was found; see also our complementary analysis in **Fig. S2a**, showing consistent results when cue reliability is treated as a categorical predictor).

### Motor prioritization strongly scales with cue reliability

Having shown that visual prioritization of the cued memory representation remained robust— and with a remarkable consistent profile—across cue-reliability conditions, we next addressed the key question: does motor prioritization (of the cued visual item’s associated action) following the cue show a similar pattern as we observed for visual prioritization?

By experimentally linking memory items’ tilt to distinct response hands, we could use patterns of EEG lateralization according to the cued item’s associated response hand to track motor prioritization in working memory. We did so by focusing on lateralization of central mu/beta (8-30 Hz) activity in electrodes C3/4 as a canonical marker of action planning (as in Kaiser et al., 2003; Pfurtscheller et al., 2005; Schneider et al., 2017; van Ede et al., 2019). Because we independently varied the cued item’s location and the response hand it required, our analysis of motor prioritization was, by construction, independent from our analyses of visual prioritization.

In contrast to visual prioritization, the amplitude of motor mu/beta lateralization profoundly scaled with cue reliability (**Fig. 2f-j**; and again, comparable mu/beta lateralization results were observed when averaging over the central electrode cluster, C3/C4, C1/2, C5/6, FC3/4, CP3/4; see **Fig. S1c-d**). We observed robust motor lateralization in the 100% and 80% reliable-cue blocks (cluster-*p* < 0.001 for both), but it became statistically absent in the 60% block (no significant cluster found). Direct contrasts revealed that mu/beta lateralization was graded by cue reliability: being significantly stronger in the 100% condition (100% vs. 80%: 391–1262 ms, cluster-*p* < 0.001; 1840–2000 ms, cluster-*p* = 0.044; 100% vs. 60%: 102–2000 ms, cluster-*p* < 0.001) as well as significantly stronger in the 80% than the 60% cue-reliability conditions (1691– 2000 ms, cluster-*p* = 0.007). Moreover, corroborating these profound modulations, for our motor prioritization signature, regression-based analysis confirmed a graded modulation by cue reliability (**Fig. 2j**; cluster-*p* < 0.001; see also our complementary analysis in **Fig. S2b**, showing consistent results when cue reliability is treated as a categorical predictor).

Thus, while visual prioritization showed no discernable modulation by cue reliability in our dataset, motor prioritization (of the response hand that was associated with the cued memory content) showed profound modulations by cue reliability. This was evident not only in the magnitude, but also in the gradual development of our motor-prioritization marker as we delineate further in the next section.

### Distinct profiles for visual vs. motor prioritization as a function of cue reliability

**Figure 2** suggested that cue reliability may not only differently affect the overall strength of our visual and motor prioritization markers, but also the gradual nature by which they develop. We finally turned to directly comparing the temporal profiles between visual and motor prioritization as a function of cue reliability. Because we could not establish a clear signature of motor prioritization in the 60% cue-reliability condition, we focused exclusively on the 100% and 80% conditions for this additional analysis.

To facilitate comparison of the temporal profiles of visual and motor prioritization, we first normalized their time courses to their respective peak value (**Fig. 3a**). A jackknife latency analysis was subsequently applied to directly compare the onset latency of visual and motor prioritization. The onset latency comparison suggested that the development of visual and motor prioritization largely overlapped in time in the 100% cue-reliability condition and did not show a significant delay (**Fig. 3a–b**, left panels; 25% peak onset latency comparison: *t*(23) = –1.259, *p* = 0.221). This is consistent with prior work showing that visual and motor prioritization can co-develop during selection from working memory following 100% reliable cues (van Ede et al., 2019; Echeverria-Altuna et al., 2025; Gresch et al., 2025). In contrast, in the 80% cue-reliability condition, our marker of visual prioritization still occurred early after the cue, but our marker of motor prioritization was now significantly delayed (**Fig. 3a–b**, right panels; 25% peak onset latency comparison: *t*(23) = –3.195, *p* = 0.004).

**Figure 3.**
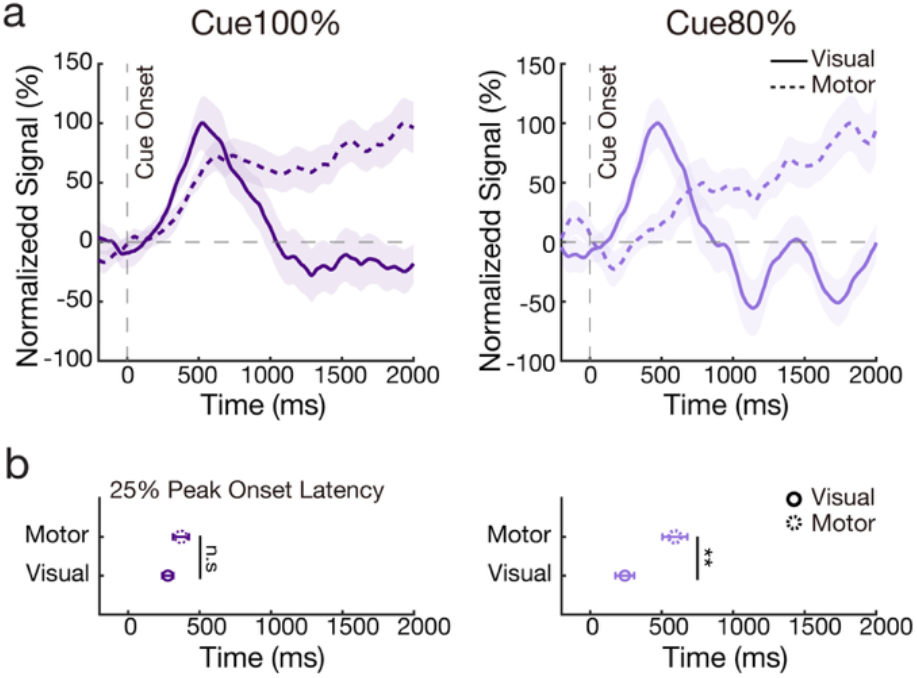
Distinct profiles for visual vs. motor prioritization as a function of cue reliability. **a)** Left panel: peak-normalized visual alpha and motor mu/beta markers following 100% reliable cues condition. Right panel: the same following 80% reliable cues condition. **b)** Left panel: onset latency (defined as the first value reaching 25% of the peak value) of visual and motor prioritization following 100% reliable cues. Right panel: the same following 80% reliable cues. Error bars represent ± 1 SEM. **, n.s represent significance level *p*<0.01, and non-significant after Bonferroni correction, respectively.

To further assess the relationship between visual and motor prioritization signals, we conducted explorative across-participant correlation analysis between the peak amplitude and onset latency (25% of peak) of alpha and mu/beta lateralization, focusing on the 100% and 80% cue-reliability conditions (as mu/beta signals were statistically absent in the 60% condition). No significant correlations were observed for peak amplitude (100%: *r* = −0.15, *p* = 0.492; 80%: *r* = 0.00, *p* = 0.992), onset latency (100%: *r* = −0.12, *p* = 0.569; 80%: *r* = 0.04, *p* = 0.884), or cross-measure relationships (all *p*s > 0.22). These null results are at least consistent with the interpretation that visual (alpha) and motor (mu/beta) prioritization can reflect dissociable processes, although we note that the absence of a significant correlation does not provide definitive evidence of independence.

We also conducted complementary analyses of alpha and mu/beta lateralization following the memory test. As shown in **Fig. S3a and S3b**, both visual and motor prioritization during retrieval were modulated by cue reliability (100%, 80%, 60%) and validity (valid vs. invalid trials), with a clearer pattern observed for the motor prioritization marker.

These additional analyses unveil how visual and motor selection do not always co-develop, but can de-couple in time (as in Echeverria-Altuna et al., 2025; Nasrawi et al., 2025), specifically following less reliable cues. When cues are less reliable, motor prioritization appears to develop more gradually (or “more carefully”), even if visual prioritization in the very same trials remains largely unaffected.

## Discussion

Our data reveal how anticipated certainty about what working-memory content will mostly likely become relevant for an upcoming task differentially modulates the proactive prioritization of visual representations and their associated manual action plans, that we jointly studied in the same task. Whereas visual prioritization remained robust and stable across cues that predicted which of two working-memory items would become tested with 60, 80 or 100% reliability, motor prioritization profoundly scaled with certainty, in strength and profile. These findings show that visual and motor prioritization in working memory are shaped by distinct mechanisms, and suggests that motor prioritization may be more cautious (and potentially more costly) in nature.

A central contribution of the present study is the demonstration that visual and motor prioritization, here studied for prioritization of working-memory content, rely on functionally distinct mechanisms that respond differently to the reliability of cues. Under fully reliable cues (100% valid), participants knew with certainty which memory item would become tested. As a result, they showed joint visual and motor prioritization (as operationalized through our EEG markers), consistent with prior work demonstrating coordinated visual–motor prioritization in working memory (van Ede et al., 2019; Boettcher et al., 2021; Nasrawi et al., 2023, 2025; Echeverria-Altuna et al., 2025; Gresch et al., 2025). However, as cue reliability decreased, visual and motor prioritization began to diverge. With moderately reliable cues (80% valid), alpha lateralization relative to the memorized location of the cued visual memory representation remained early and robust – consistent with prior findings that posterior alpha lateralization reliably indexes visual prioritization in working memory even under uncertainty (Gunseli et al., 2015; Myers et al., 2015; Foster et al., 2017; Fu et al., 2022; van Ede and Nobre, 2023; Wang and van Ede, 2024). In contrast, under lower certainty, central mu/beta dynamics related to the prospective response hand were profoundly weakened (and developed more gradually). This suggests greater susceptibility of motor prioritization to certainty (cf. Tzagarakis et al., 2010, 2015, 2021; Nasrawi and van Ede, 2022; Nasrawi et al., 2025). This dissociation became even more pronounced under low cue certainty (60% valid) where motor-related mu/beta lateralization was no longer evident, despite clearly preserved visual prioritization in the same trials. Such de-coupling between visual and motor prioritization aligns with recent work using complementary manipulations that corroborate the notion that visual and motor prioritization can be dissociated, such as under variations in memory load (Nasrawi et al., 2025) or temporal expectations (Echeverria-Altuna et al., 2025).

Our data suggests that, under uncertainty, humans are more willing to commit to visual prioritization (that may prove wrong) than to motor prioritization (that may prove wrong). We speculate that under lower certainty, participants perhaps prefer to plan for multiple potential actions in parallel (Cisek, 2007; Gallivan and Wood, 2009; though note how the parallel-planning hypothesis has also been questioned: Wong and Haith, 2017; Alhussein and Smith, 2021) rather than to commit to the prioritization of specific actions. Strikingly, however, this putative strategy does not appear to apply similarly to the commitment to prioritize visual representation under the exact same degree of certainty. Possibly, it is less costly to have to later re-prioritize another visual representation upon an unexpected memory test (cf. Wang and van Ede, 2024), than it may be to have to re-prioritize an alternative course of action. Future studies are required to corroborate this assertion.

Our findings connect to ample prior studied that reported how posterior alpha lateralization scales with cue reliability, in both perception and working memory (Poch et al., 2014, 2017; Schneider et al., 2015, 2016; Wolff et al., 2017; van Ede, 2018; Günseli et al., 2019; Fu et al., 2022; Macedo-Pascual et al., 2022; Li et al., 2023; Wang and van Ede, 2025a). While we observed a similarly graded trend with numerically stronger lateralization following more reliable cues, this trend did not reach significance in our current dataset. Possibly this is owing to the nature of our task, in which participants could prioritize not only visual representations but also manual action plans, which was not the case in any of the aforementioned studies. Possibly, in such a task, the graded use of the cue may shift toward the motor prioritization component. Indeed, in our data, central mu/beta lateralization weakened and developed more gradually as cue uncertainty increased. At the same time, we note how it is not the case that because of the nature of our task, participants skipped visual prioritization altogether. In fact, we observed highly robust alpha lateralization as a signature of visual prioritization in our task, even if, unlike our marker of motor prioritization, it showed only modest, and non-significant, modulations by cue reliability. We note that the clear modulation by cue reliability on the motor-prioritization marker – measured within the same participants, task, and trials as the visual-prioritization marker – indicates that the overall design and dataset were capable of yielding robust cue-reliability effects. This provides relevant context for interpreting the comparatively weaker and non-significant cue-reliability effects on our visual-prioritization marker, though we note that this may also reflect limited sensitivity of this particular visual-prioritization marker.

In the current study, we operationalized visual and motor prioritization through posterior alpha and central mu/beta lateralization, building on (van Ede et al., 2019; Boettcher et al., 2021; Nasrawi et al., 2023, 2025; Echeverria-Altuna et al., 2025; Gresch et al., 2025). This revealed distinct patterns of visual and motor prioritization as a function of cue reliability. At the same time, it is important to acknowledge that these two human-neuroscience spectral EEG markers are but two specific neural markers for such processes and unlikely capture visual and motor prioritization exhaustively. Accordingly, we do not take our reported lack of action prioritization following 60% reliable cues to imply that no action prioritization took place at all, as other markers may still show such prioritization. Even so, with these markers at our disposal, we established robust motor prioritization in the 80% and 100% conditions, confirming overall suitability. We further acknowledge that our markers may entail multiple underlying mechanisms (putatively reflecting a change in the excitability across relevant visual and motor areas; Engel and Fries, 2010; Jensen and Mazaheri, 2010; Spitzer and Haegens, 2017; Iemi et al., 2022), and that what we have referred to as “prioritization” likely incorporates multiple processes, including visual and motor selection, and subsequent motor preparation.

One important consideration is whether the observed modulation of mu/beta lateralization reflects purely central motor preparation or is partly driven by sub-threshold motor commands at the peripheral level. Previous work has shown that subtle EMG activity can track decision-related processes prior to overt movement (e.g., Selen et al., 2012; Papaioannou and Dimitriou, 2021; Rungta and Murthy, 2023), raising the possibility that covert muscle activation may contribute to our preparatory EEG signals. Although our task required no overt response during the cue–delay interval, and prior studies suggest that lateralized mu/beta activity reflects central motor preparation in similar paradigms (e.g., Schneider et al., 2017; van Ede et al., 2019; Boettcher et al., 2021), we cannot fully exclude potential contributions from sub-threshold motor activity. Yet, even if subthreshold motor activity in the periphery may contribute to our EEG marker, our marker would nonetheless signal prioritization of the relevant action linked to specific working-memory content. While addressing this possibility would be valuable for refining what our motor prioritization marker reflects, we note that whether or not peripheral signals contribute to our motor marker in our task does not fundamentally change our main conclusion that cue reliability differentially modulates visual and motor prioritization during and/or after selection from working memory.

Notwithstanding these drawbacks, these spectral markers of lateralization in the EEG showed differential sensitivity to cue reliability, even when studied in the same participants, in the same task, and in the same trials. Our data thus suggests that the certainty about anticipated task demands differentially shapes the proactive prioritization of relevant visual representations and their associated manual action plans, whereby the prioritization of action plans appears to be more cautious in nature.

## Acknowledgements

This work was supported by an ERC Starting Grant from the European Research Council (MEMTICIPATION, 850636) and an NWO Vidi Grant from the Dutch Research Council (14721) to F.v.E. We also thank Narges Naghibi for her assistance with data collection.

## Supplementary Materials

**Figure S1.**
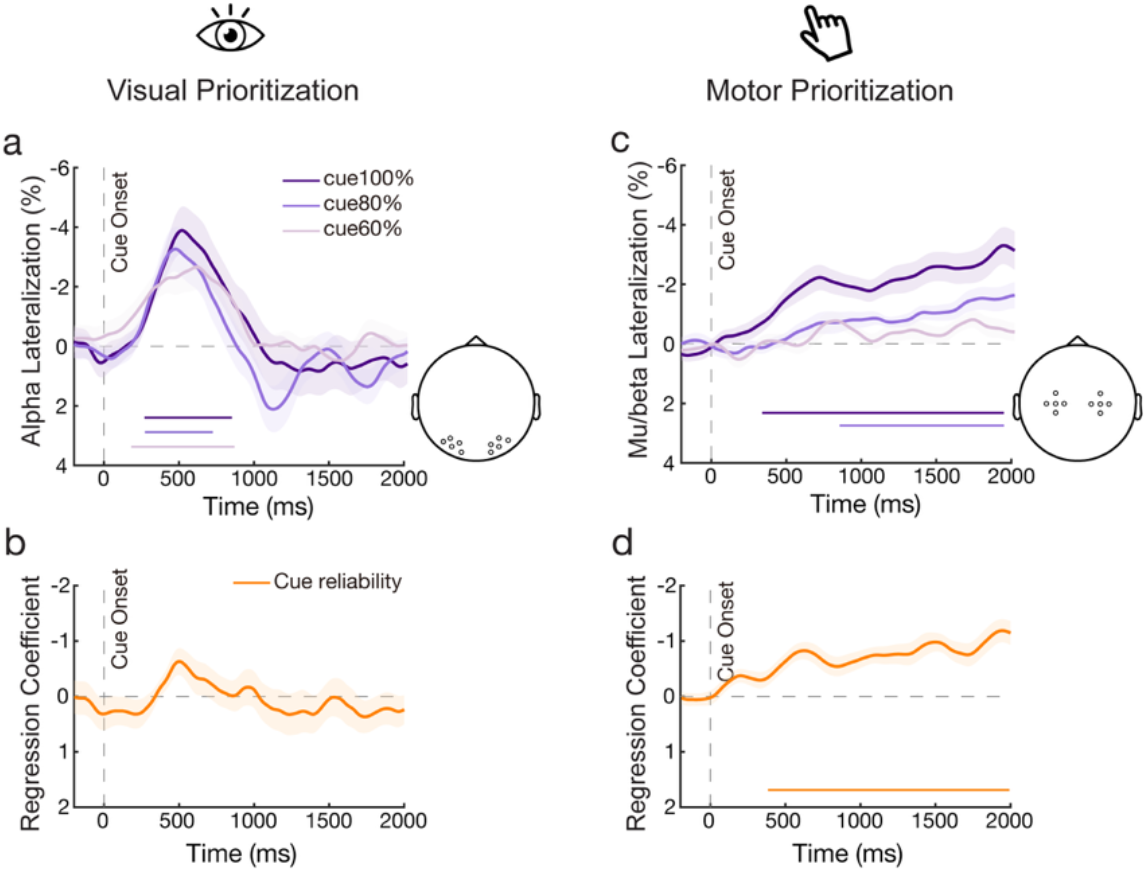
Cue reliability diEerentially modulates proactive visual and motor prioritization during working memory also when using electrode clusters. **a)**. Time courses of alpha lateralization (8-12 Hz) relative to the cued item’s memorized location in posterior electrodes cluster (PO7/8, PO3/4, P7/P8, P5/6, O1/O2) across three levels of cue reliability. **b)** The group-level average of beta estimates from general linear model fitting trial-wise alpha lateralization (8-12 Hz) with cue-reliability as a continuous predictor. **c-d)** Similar as **a-b**, but for 8-30 Hz motor lateralization in central electrodes cluster (C3/C4, C1/2, C5/6, FC3/4, CP3/4) relative to the prospective response hand associated with the cued item. Colored horizontal lines above the x-axes indicate significant clusters (p<0.05). Shading and error bars represent ± 1 SEM, n = 24.

**Figure S2.**
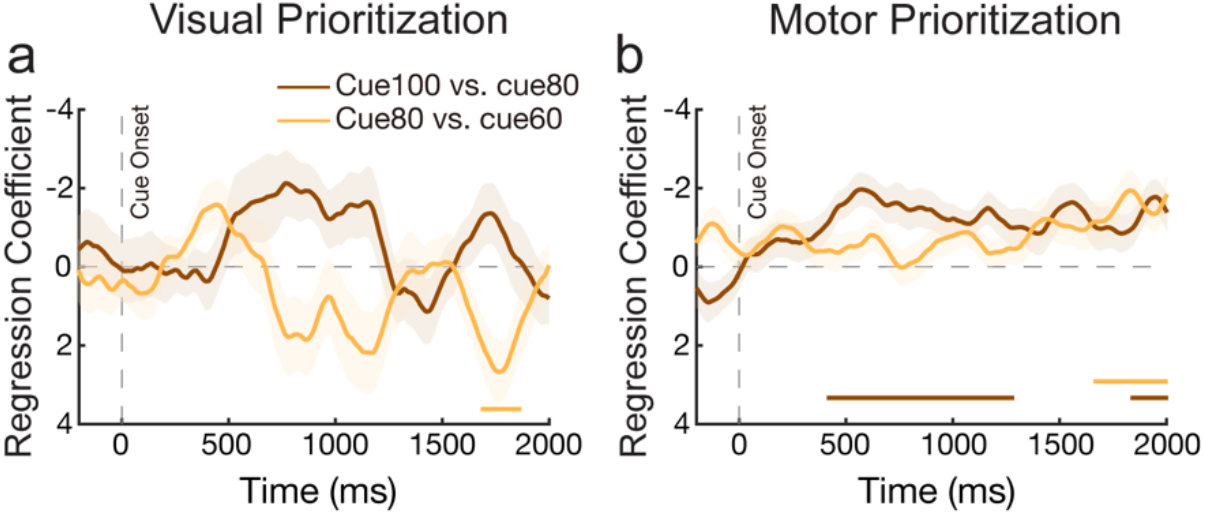
Cue reliability (treated as a categorical predictor) differentially predicts visual and motor prioritization. **a)** Group-level averages of beta estimates from a general linear model fitting trial-wise alpha lateralization (8–12 Hz), with cue reliability treated as a categorical predictor (high vs. medium vs. low). The 80% cue-reliability condition was used as the baseline reference. A permutation cluster test revealed a small late significant cluster for the 80% vs. 60% comparison (cluster-*p* = 0.030, 1648–1852 ms after cue onset). **b)** Same as in **a**, but for mu/beta lateralization. Significant clusters were observed for the 80% vs. 100% comparison (cluster-*p* < 0.001, 391–1262 ms; cluster-*p* = 0.049, 1840– 2000 ms after cue onset), and for the 80% vs. 60% comparison (cluster-*p* = 0.010, 1691–2000 ms after cue onset). Colored horizontal bars above the x-axes indicate significant clusters (*p* < 0.05). Shading and error bars represent ±1 SEM; n = 24.

**Figure S3.**
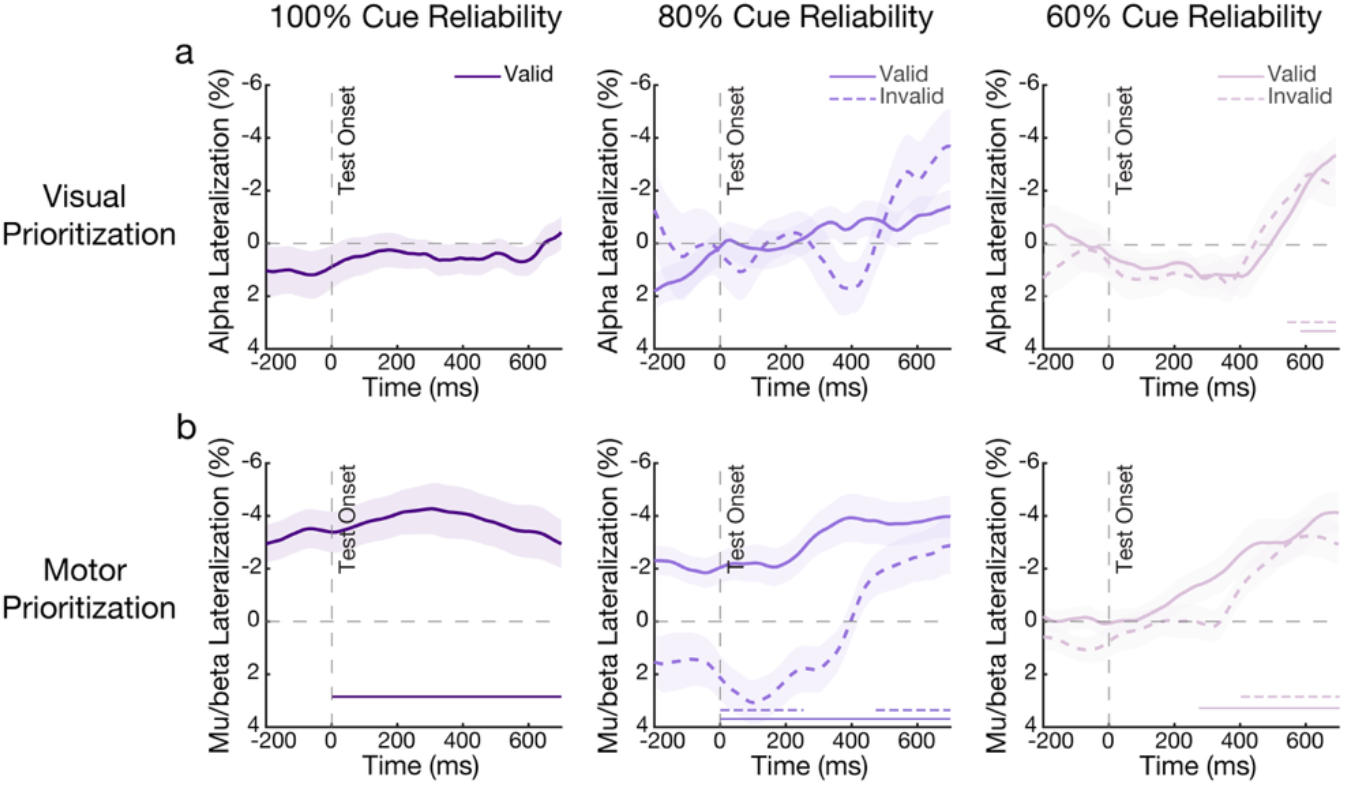
Visual and motor selection following the working-memory test as a function of cue reliability and validity. **a)** Time courses of alpha lateralization (8–12 Hz), relative to the memorized location of the tested item, measured at the posterior electrode pair (PO7/8) across three levels of cue reliability, separately for valid and invalid trials following test onset. Permutation cluster tests revealed significant alpha lateralization during memory retrieval in the 60% cue-reliability condition for both valid and invalid trials (valid: cluster *p* = 0.021, 590–700 ms; invalid: cluster *p* = 0.030, 559–700 ms post-test onset). **b)** Same as in (**a**), but for mu/beta lateralization (8–30 Hz) relative to the response hand associated with the tested item, measured at the central electrode pair (C3/C4). Significant clusters were observed in the 100% condition (cluster *p* < 0.001, 0–700 ms), in the 80% valid condition (cluster *p* < 0.001, 0–700 ms), and in the 80% invalid condition, which showed a reversal pattern (positive cluster: *p* = 0.006, 0–238 ms; negative cluster: *p* = 0.014, 461–700 ms). In the 60% condition, significant clusters were found for both valid (cluster *p* < 0.001, 262–700 ms) and invalid trials (cluster *p* < 0.001, 0–700 ms). Colored horizontal bars above the x-axes indicate significant clusters (*p* < 0.05). Shaded areas represent ±1 SEM (n = 24).

Figure S3 shows alpha and mu/beta lateralization following the memory test. Both visual and motor prioritization during retrieval were modulated by cue reliability (100%, 80%, 60%) and validity (valid vs. invalid trials), with a clearer pattern observed for our motor prioritization marker. In the 100% condition, strong motor prioritization persisted from the cue through test onset (**Fig. S3b**, left panel). In the 80% condition, where valid-cue trials were more likely, participants appeared to ‘flip’ their motor prioritization on invalid-cue trials after test onset—an effect not observed for valid trials (**Fig. S3b**, middle panel). A similar but non-significant trend was observed in visual selection (**Fig. S3a**, middle panel). This pattern is consistent with the observed motor prioritization following the cue, suggesting that participants initially prepare responses based on the cue and subsequently adjust their motor plans when the cue proves invalid.

In contrast, in the 60% condition, participants prioritized the required response hand at test onset regardless of cue validity (**Fig. S3a,b**, right panels). This aligns with the reported lack of motor prioritization following the cue, indicating that participants defer response preparation until the test display when cue reliability is low.

Together, these findings suggest a dynamic adjustment of visual and motor prioritization, and corroborate our primary findings from the cue-locked stage.

